# Frequency-specific coactivation patterns in resting-state and their alterations in schizophrenia: an fMRI study

**DOI:** 10.1101/2021.07.04.451042

**Authors:** Hang Yang, Hong Zhang, Xin Di, Shuai Wang, Chun Meng, Lin Tian, Bharat Biswal

**Author notes:** **Correspondence:** Dr. Chun Meng, The Clinical Hospital of Chengdu Brain Science Institute, MOE Key Laboratory for Neuroinformation, Center for Information in Medicine, School of Life Science and Technology, University of Electronic Science and Technology of China, No. 2006, Xiyuan Avenue, Chengdu 611731, China;, Dr. Lin Tian, Department of Psychiatry, The Affiliated Wuxi Mental Health Center of Nanjing Medical University, No. 156 Qianhu Road, Binhu District, Wuxi 214151, China;, Dr. Bharat B. Biswal, 607 Fenster Hall, University Height, Newark, NJ, 07102, USA.

## Abstract

The resting-state human brain is a dynamic system that shows frequency-specific characteristics. Coactivation pattern (CAP) analysis has been recently used to identify recurring brain states sharing similar coactivation configurations. However, whether and how CAPs differ across different sub-frequency bands are unknown. In the current study, in addition to the typical low-frequency range (0.01 - 0.08 Hz), the spatial and temporal characteristics of CAPs in four sub-frequency bands, slow-5 (0.01 - 0.027 Hz), slow-4 (0.027 - 0.073 Hz), slow-3 (0.073 - 0.198 Hz), and slow-2 (0.198 - 0.25 Hz), were studied. Six CAP states were obtained for each band., The CAPs from the typical frequency range were spatially largely overlapped with those in slow-5, slow-4 and slow-3 but not with those in slow-2. With the increase of frequency, the CAP state became more unstable and resulted in an overall shorter persistence. The spatial and temporal characteristics of slow-4 and slow-5 were further compared, because they constitute most power of the resting-state fMRI signals. In general, slow-4 showed stronger coactivations or co-deactivations in subcortical regions, while slow-5 showed stronger coactivations or co-deactivations in large-scale cortical networks such as the dorsal attention network. Lastly, frequency-dependent dynamic alterations were also observed in schizophrenia patients. Combining the information obtained from both slow-5 and slow-4 increased the classification accuracy of schizophrenia patients than only using the typical range. In conclusion, our results revealed that the spatial and temporal characteristics of CAP state varied at different frequency bands, which could be helpful for identifying brain alterations in schizophrenia.

## 1. Introduction

The human brain is a dynamic system, and the resting-state functional connectivity (RSFC) has been proved to be temporally varied (Chang and Glover 2010). The conventional dynamic functional connectivity (dFC) approach segments the time-series using sliding-window and calculates the interregional Pearson correlation within each window (Chang and Glover 2010; Hutchison et al. 2013). Moreover, recurring connectivity configurations across windows could be grouped as FC-states (Allen et al. 2014). These FC-states were found to be related to cognitive and physiological states such as vigilance (Wang, Ong, et al. 2016), self‐generated thought (Marusak et al. 2017), eyes open and closed (Weng et al. 2020), and also disease alterations (Guo et al. 2019; Li, Dong, et al. 2020; Damaraju et al. 2014). However, the choice of window length and window shape remained to be optimized (Zalesky and Breakspear 2015; Shakil, Lee, and Keilholz 2016), and the temporal resolution is also relatively low as the recommended window length is about 30-50 seconds (Hutchison et al. 2013).

Instead of estimating brain states using the sliding-window dFC maps, brain states can also be identified based on recurring coactivation patterns (CAPs) from each single frame (Liu and Duyn 2013). The CAP analysis was first performed using a seed-based approach and a threshold was needed to select the suprathreshold frames (Liu and Duyn 2013), then it was extended to a seed-and-threshold-free approach (Liu, Chang, and Duyn 2013). Comparing with sliding-window dFC maps, CAPs are more direct measurements of brain activities without any statistic or mathematic calculation. It also has a better temporal resolution and does not require predefined parameters such as window length, as the analytical unit of CAP analysis is a single volume. Besides, our previous study has shown the robustness of CAPs across several technique flexibilities and independent cohorts, and reproducible alterations were also obtained between schizophrenia patients and healthy controls (Yang et al. 2021). Recently, CAP analysis has been used to study the altered brain dynamics in patients with depression (Kaiser et al. 2019) and Alzheimer’s disease (Ma et al. 2020). In addition, Li and colleagues concatenated a set of task activation maps from the Human Connectome Project, and they identified robust anti-correlated functional networks (default network) across multiple tasks (Li, Dahmani, et al. 2020), suggesting that CAP analysis could also be utilized in task fMRI.

Besides the temporal dynamics contained in the resting-state fMRI blood oxygen level dependent (BOLD) signals, frequency-dependent information also exists. Previous resting-state fMRI studies mainly focused on the low-frequency oscillation (LFO) which fluctuates at the typical low-frequency band (0.01 – 0.08/0.1 Hz), as the LFO is thought to reflect the intrinsic neuronal fluctuations (Biswal et al. 1995). Although the higher frequency fluctuations are regarded as physiological noise such as respiration-induced and cardiac noise (Cordes et al. 2001), RSFC above 0.1 Hz and the potential physiological significance of high-frequency BOLD signal (Chen and Glover 2015) is under debate. To measure the effects of neural activity in different frequency bands, the frequency range was generally subdivided into four sub-frequency bands, including slow-5 (0.01 - 0.027 Hz), slow-4 (0.027 - 0.073 Hz), slow-3 (0.073 - 0.198 Hz) and slow-2 (0.198 - 0.25 Hz) based on previous electrophysiological (Buzsaki and Draguhn 2004) and fMRI studies (Zuo et al. 2010). Inhomogeneous spatial amplitude of low-frequency fluctuations (ALFF) distribution between slow-4 and slow-5 were observed (Zuo et al. 2010). Furthermore, frequency-specific ALFF changes have been found in disease groups such as mild cognitive impairment (Han et al. 2011), Parkinson’s disease (Hou et al. 2014) and depression (Wang, Kong, et al. 2016) between slow-4 and slow-5. Besides, frequency-specific effects have also been widely reported in functional connectivity (Gohel and Biswal 2015), regional homogeneity (ReHo) (Yu et al. 2016), and brain networks (Xue et al. 2014). These findings indicate the underlying frequency-dependent brain activity and frequency-specific disease alterations. While for CAPs, whether and how would the spatial and temporal characteristics change with the frequency band is unknown and remains to be explored.

Schizophrenia is a mental disorder with globally altered brain functions, and the aberrant brain dynamics found in schizophrenia patients (SZ) have the potential to be the biomarker to reveal the complex pathology of this disease (Du et al. 2017; Kottaram et al. 2018). The abnormal dynamic brain graphs (Yu et al. 2015) and network reconfigurations have also been identified in SZ (Reinen et al. 2018). Our previous study has shown the altered CAP state dynamics of SZ patients were not only involved with the triple-network, but also extend to other primary and high-order networks (Yang et al. 2021). Besides the dynamic brain changes, SZ patients also showed frequency-specific alterations in several aspects, including ALFF (Yu et al. 2014; Gohel et al. 2018; Meda et al. 2015; Hare et al. 2017), ReHo (Yu et al. 2013), BOLD variability (Zhang, Yang, and Cai 2020), as well as functional connectivity (Wang et al. 2017; Han et al. 2017). Furthermore, Zou and colleagues distinguished schizophrenia patients from healthy controls using the dFC estimated at different frequency bands (Zou and Yang 2019), and Luo et al. found that SZ showed distinct dFC strength alterations in slow-4 and slow-5 (Luo, He, et al. 2020), these results suggest the underlying frequency-specific dynamic alterations in psychosis. Based on the above findings, SZ patients may also be affected by frequency-specific CAP alterations that need further investigations.

The purpose of this study is to test whether the frequency-dependent effects can be observed using coactivation patterns. Specifically, the typical range (0.01 – 0.08 Hz) and four sub-frequency bands from slow-5 to slow-2 were analyzed, and CAP analysis was performed in each frequency band separately. Then, the spatial and temporal characteristics varied with frequency bands were evaluated, and particularly the results of slow-4 and slow-5 were statistically compared, as these two sub-frequency bands are within the typical low-frequency range and have been widely studied in previous studies. Finally, the frequency-dependent CAPs were applied to schizophrenia patients, and the frequency-specific disease alterations were studied in this work.

## 2. Materials and methods

### 2.1 Participants

All participants were scanned at the Department of Medical Imaging, Wuxi People's Hospital, Nanjing Medical University. Four subjects were excluded due to the large headmotion. As shown in Table 1, 69 schizophrenia patients (35 males/34 females, 46.06 ± 10.96 years) and 97 healthy controls (56 males/41 females, 40.36 ± 14.77 years) remained for the current study. Positive and Negative Syndrome Scale (PANSS) was performed for all schizophrenia patients to evaluate their symptom severity. This research was approved by the Medical Ethics Committee of Wuxi Mental Health Center, Nanjing Medical University (study number: WXMHCIRB2012LLKY001), and was conducted in accordance with the Declaration of Helsinki guidelines. The written informed consent was obtained from all participants.

**Table 1.** The demographic information for the WuXi cohort

| WuXi | All - HC<br>(n = 97) | Matched - HC<br>(n = 69) | SZ<br>(n = 69) | P value |
| --- | --- | --- | --- | --- |
| Age | $40.36 \pm 14.77$ | $45.84 \pm 11.89$ | $46.06 \pm 10.96$ | 0.9112<sup>a)</sup> |
| Gender (M \ F) | 56 \ 41 | 35 \ 34 | 35 \ 34 | 1<sup>b)</sup> |
| Disease duration | - | - | $19.84 \pm 10.96$ | - |
| PANSS positive | - | - | $20.06 \pm 4.59$ | - |
| PANSS negative | - | - | $23.78 \pm 3.84$ | - |
| PANSS general | - | - | $41.67 \pm 5.27$ | - |
| PANSS total | - | - | $85.51 \pm 9.50$ | - |

Data are expressed as mean ± SD (SD: standard deviation).
Abbreviations: PANSS, Positive and Negative Syndrome Scale.
<sup>a)</sup> two-sample t-test; <sup>b)</sup> chi-square cross-table test.

### 2.2 fMRI Data Acquisition

Three-dimensional T1-weighted images and resting-state fMRI scans were collected using a 3.0 T Magnetom TIM Trio (Siemens Medical System). Structural MRI images were acquired using a 3D-MPRAGE sequence with the following parameters: TR/TE = 2530/3.44 ms, flip angle = 7°, FOV = 256 mm, matrix size = 256 × 256, voxel size = 1 × 1 × 1 mm^3^, slice thickness = 1 mm and slice number = 192. Resting-state fMRI data were obtained using a single-shot gradient-echo echo-planar-imaging sequence, with TR/TE = 2000/30 ms, flip angle = 90°, FOV = 220 mm, matrix size = 64 × 64, voxel size = 3.4 × 3.4 × 4 mm^3^, slice thickness = 4 mm, slice number = 33, and 240 volumes were collected for each subject.

### 2.3 Data Preprocessing

All structural and functional MRI images were preprocessed using DPABI (http://rfmri.org/dpabi). The T1-weighted images were first coregistered to the functional images, and then segmented into gray matter, white matter and cerebrospinal fluid by using DARTEL. For the resting-state fMRI images, the first 5 time points were removed to avoid instability of the scanner, and the remaining images were realigned to correct the head movement. Framewise displacement (FD) was calculated for each subject (Di and Biswal 2015), and subjects with maximum translation or rotation FD greater than 2 mm or 2° were excluded from further analysis. The fMRI images were normalized to the Montreal Neurologic Institute (MNI) space using the deformation field maps obtained from the T1 segmentation, and resampled to 3 × 3 × 3 mm^3^. The mean white matter, cerebrospinal fluid and global signal, and 24 head motion parameters (Friston et al. 1996) were regressed from the time series. The time series was further detrended and temporal filtered. Besides the typical filtering bandpass (0.01 - 0.08 Hz), another four sub-bands including slow-5 (0.01 - 0.027 Hz), slow-4 (0.027 - 0.073 Hz), slow-3 (0.073 - 0.198 Hz) and slow-2 (0.198 - 0.25 Hz) were employed separately based on the previous study (Zuo et al. 2010). Finally, all images were smoothed using an 8 mm FWHM Gaussian kernel.

Similar to our previous study, the mean BOLD time series was extracted from 408 ROIs separately (Yang et al. 2021), which includes 400 cortical regions (Schaefer et al. 2018) and 8 subcortical regions (bilateral caudate nucleus, putamen, globus pallidus and amygdala) from the AAL template (Tzourio-Mazoyer et al. 2002). The 400 cortical regions belong to 7 networks, including the visual network (VN), somatomotor network (SMN), dorsal attention network (DAN), ventral attention network (VAN), limbic network, fronto-parietal network (FPN) and default mode network (DMN).

### 2.4 Coactivation Pattern Analysis

The coactivation pattern (CAP) analysis is a data-driven method based on the k-means clustering, and it is supposed to identify recurring whole-brain coactivation states. The same analysis pipeline from our previous work (Yang et al. 2021) was used to detect the CAP states in different frequency bands.

In brief, there were 235 volumes for each subject, and each volume was characterized by the activation level of 408 ROIs. The time series of each ROI was first normalized using z-score independently, and the absolute value of Z indicates the activation deviation from its baseline. Then, K-means clustering was performed based on all volumes from the 97 HC subjects, and volumes sharing similar coactivation profiles were grouped into the same CAP state. The spatial map of each CAP was obtained by averaging across volumes belonging to the state, and divided by their standard deviation to generate a Z-map (Liu and Duyn 2013). Pearson correlation was used to measure the spatial similarity between volumes and CAP states. As for the SZ subjects, their volumes were assigned to the obtained CAP state with the highest spatial similarity. The cluster number K was tested from 2 to 21, and the silhouette score (Rousseeuw 1987) was used to determine the cluster number. Our previous work identified six robust CAP states in the typical range (H. Yang et al., 2021). We found six clusters were also suitable for the four sub-frequency bands, and their silhouette score curves were shown in Supplementary Figure S1.

### 2.5 CAP state temporal dynamic measures

The temporal dynamic properties among the six CAP states were evaluated using four CAP metrics at the individual level. **Fraction of time** represents the proportion of time occupied by one state. **Persistence** describes the average dwell time, and **Counts** records the frequency of one state that occurs across the scan. In addition to these state dominances that capture the inner-state dynamics, the **transition probability** between states was also measured and presented in the supplementary materials.

### 2.6 Statistical Analysis

For comparisons within the HC group, all 97 HC subjects were analyzed, and for comparisons between SZ and HC, only age- and gender-matched HC subjects were included (69 HC and 69 SZ). For the demographic data, two-sample t-test was used to compare the age difference between SZ and HC, and chi-square cross-table test was used to test their gender difference.

In this study, the spatial and temporal characteristics of CAPs in slow-5 (0.01 - 0.027 Hz) and slow-4 (0.027 - 0.073 Hz) were further compared, as they constitute most power of the typical low-frequency range (0.01 – 0.08 Hz). For the CAP results within the HC group, the six CAPs of slow-5 and slow-4 were compared at the ROI level using paired t-test, and Bonferroni correction was used to correct the multiple comparisons (p < 0.05/408) for each state. To better illustrate the 408 ROIs’ group averaged activation level in slow-4 and slow-5 (the first two columns in Figure 4), boxplots were plotted for the six CAPs, and the 408 ROIs were categorized into the seven networks. As for the temporal dynamic measures, the CAP matrices were compared between slow-5 and slow-4 using paired t-test, and p values were false-discovery rate (FDR) adjusted. Besides, the between-state temporal differences were also examined using paired t-test in slow-4 and slow-5 separately, and Bonferroni correction was performed.

Furthermore, the CAP dynamic differences between SZ and HC in slow-5 and slow-4 were studied, and the group × frequency interaction effects were estimated using a two-way repeated-measures analysis of variance (ANOVA) with age and gender as covariates, and FDR correction was performed to account for the multiple comparisons. For post hoc comparisons, two-sample t-test (with age and gender controlled) was performed to clarify the group differences, and paired t-test was used to detect the frequency effects. FDR correction was performed across all post hoc tests.

### 2.7 Classification Analysis

To further investigate that whether combining slow-5 and slow-4 contains more information and improves the classification accuracy of schizophrenia patients than just using the typical range, eight classification models have been built using temporal or spatial features from typical range, slow-5, slow-4 separately, and combined slow-5 and slow-4. In this study, we used a similar classification model and feature selection strategy with our previous work (Yang et al. 2020), and the details were illustrated in the supplementary materials.

## 3. Results

### 3.1 Spatial and temporal properties of CAPs at different frequency bands

The CAP analysis was performed in all 97 HC subjects in the typical range and four sub-frequency bands (slow-5 to slow-2) separately. As shown in Figure 1, typical intrinsic high-order (e.g., FPN and DMN) and primary networks (e.g., VN and SMN) can be observed in the typical range, slow-5, slow-4 and slow-3. While some of them, particularly the high-order networks disappeared in slow-2, for instance, the DMN and FPN cannot be found in any state in slow-2. Pearson correlation was calculated between each pair of states to quantify the CAPs spatial similarities between the typical range and the other four frequency bands (Figure 2). All six CAP states showed high spatial one-to-one correspondence between slow-4 and the typical range, as can be observed from the diagonal of the matrix (Figure 2B), followed by slow-5 and slow-3. As for slow-2, only State 2 and State 3 were similar to the typical range.

**Figure 1.**
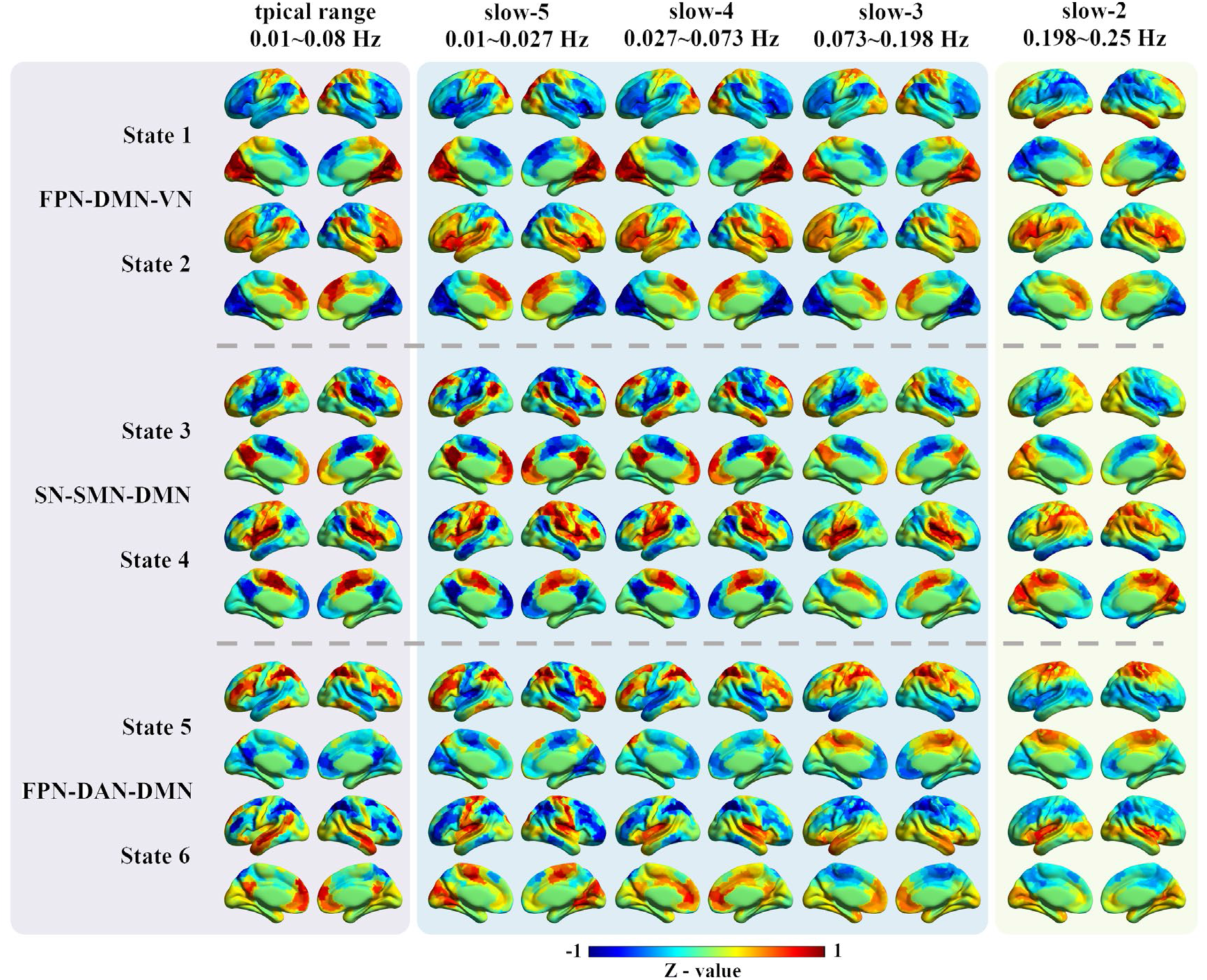
The spatial patterns for the six CAP states in different frequency bands. The first column shows the six CAP states in the typical range, and the six states were grouped into three pairs with opposite coactivation profiles. The following four columns show the six CAP states from slow-5 to slow-2. For each ROI, the Z-value means the degree of activation deviation from its baseline. The warm color indicates a relatively stronger BOLD response than its baseline amplitude, and vice versa for the cold color. Abbreviations: DAN, dorsal attention network; DMN, default mode network; FPN, fronto-parietal network; SN, salience network; SMN, somatomotor network.

**Figure 2.**
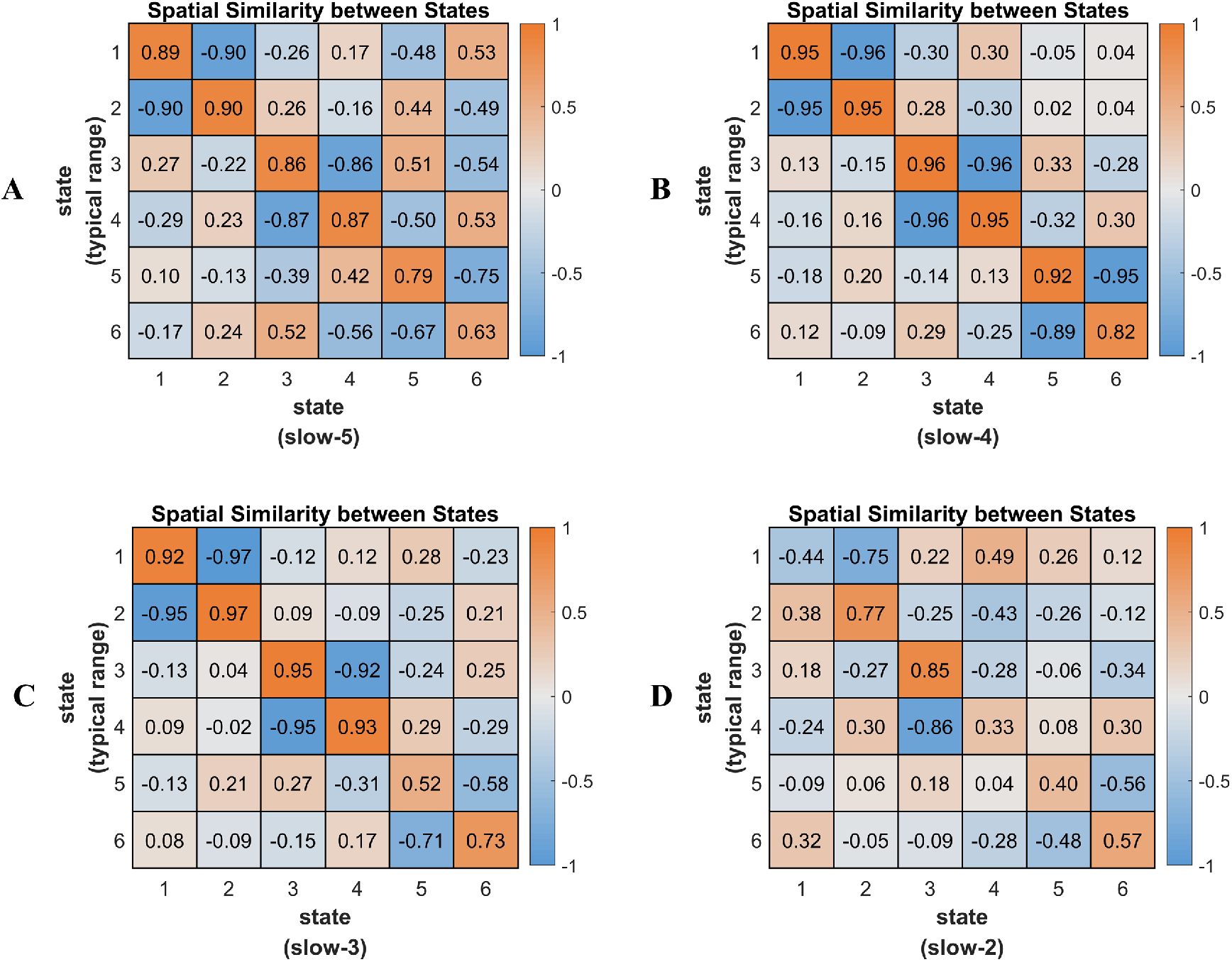
The CAP spatial similarity between the typical range and four sub-frequency bands. Pearson correlation was calculated to measure their spatial similarity, and the colorbar shows the R-value.

The absolute value of activation amplitude of each ROI indicates the deviation from its baseline activation level (Z value = 0), and was defined as activation deviation in this work. A larger activation deviation means a stronger positive activation or stronger negative deactivation. For example, compared with other brain areas, regions within the visual network exhibited stronger positive activation in State 1, and stronger negative deactivation in State 2. Hence, we said that State 1 and State 2 showed larger activation deviation in the visual network, and the visual network was the dominant network for State 1 and State 2.

In our previous study, the six CAP states were grouped into three pairs (State 1 and State 2, State 3 and State 4, State 5 and State 6) in the typical range, and the paired CAP states were characterized by opposite coactivation profiles. In the typical range, State 1 and State 2 were mainly dominated by VN, FPN and DMN, State 3 and State 4 were mainly dominated by SN, SMN and DMN, and State 5 and State 6 were mainly dominated by FPN, DAN and DMN. The between-state spatial similarity matrix was measured for each sub-frequency band independently in this study, and the spatially opposite CAP pairs can also be observed in the four sub-frequency bands (supplementary Figure S2). For instance, State 3 and State 4 belong to an opposite CAP-pair. The DMN was activated, and the SMN and SN were deactivated in State 3, while the DMN was deactivated, and the SMN and SN were activated in State 4.

The temporal dynamics in the typical range and four sub-frequency bands were then compared and shown in Figure 3. The mean fraction of time was comparable across the six CAP states for all frequency bands, around 15% to 20%. As for the persistence, with the decrease of frequency bands from slow-2 to slow-5, each state persisted longer before it transfers to another state. For slow-2 and slow-3, each state would only persist for about 2 seconds, and increased to about 5 seconds for slow-4 and 12 seconds for slow-5. The persistence of the typical range was between slow-4 and slow-5, which was around 6 seconds. On the contrary, counts (occurrences of state) decreased with the decrease of the frequency band.

**Figure 3.**
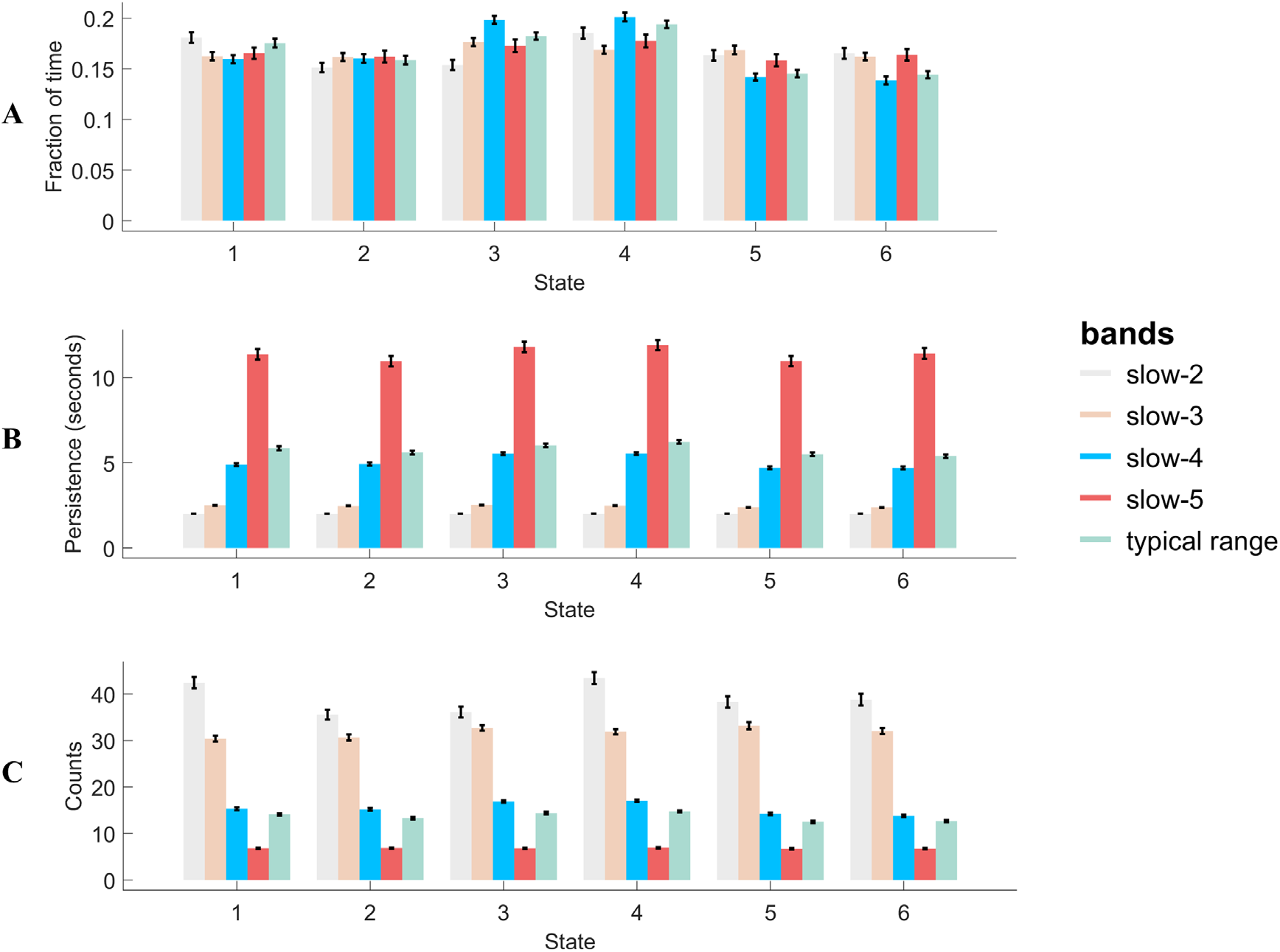
The state temporal dominances (fraction of time, persistence and counts) in the typical range and four sub-frequency bands. The error bar shows the standard error.

### 3.2 Specific spatial and temporal characteristics of CAPs in slow-4 and slow-5

To further investigate that, within the typical low-frequency range (0.01 - 0.08 Hz), whether the two popular studied sub-frequency bands (slow-4 and slow-5) showed frequency-specific spatial and temporal characteristics, the spatial maps and CAP dynamics were statistically compared within the HC group. As described in the supplementary Figure S5, for the six CAP states between slow-4 and slow-5, one-to-one correspondence can be established based on CAPs’ spatial similarity. As the six states were grouped into three pairs with opposite coactivation patterns (State 1 and 2, State 3 and 4, State 5 and 6), similar regions showed frequency-specific activation differences for the paired states, hence we only showed half of the results (Figure 4), and the remained results were described in the supplementary Figure S6. To make the naming rule consistent with our previous results based on the typical range (Figure 1), the three states were still named as FPN-DMN-VN, SN-SMN-DMN and FPN-DAN-DMN. The group averaged activation levels of the seven networks in slow-4 and slow-5 were also shown in the last column of Figure 4.

**Figure 4.**
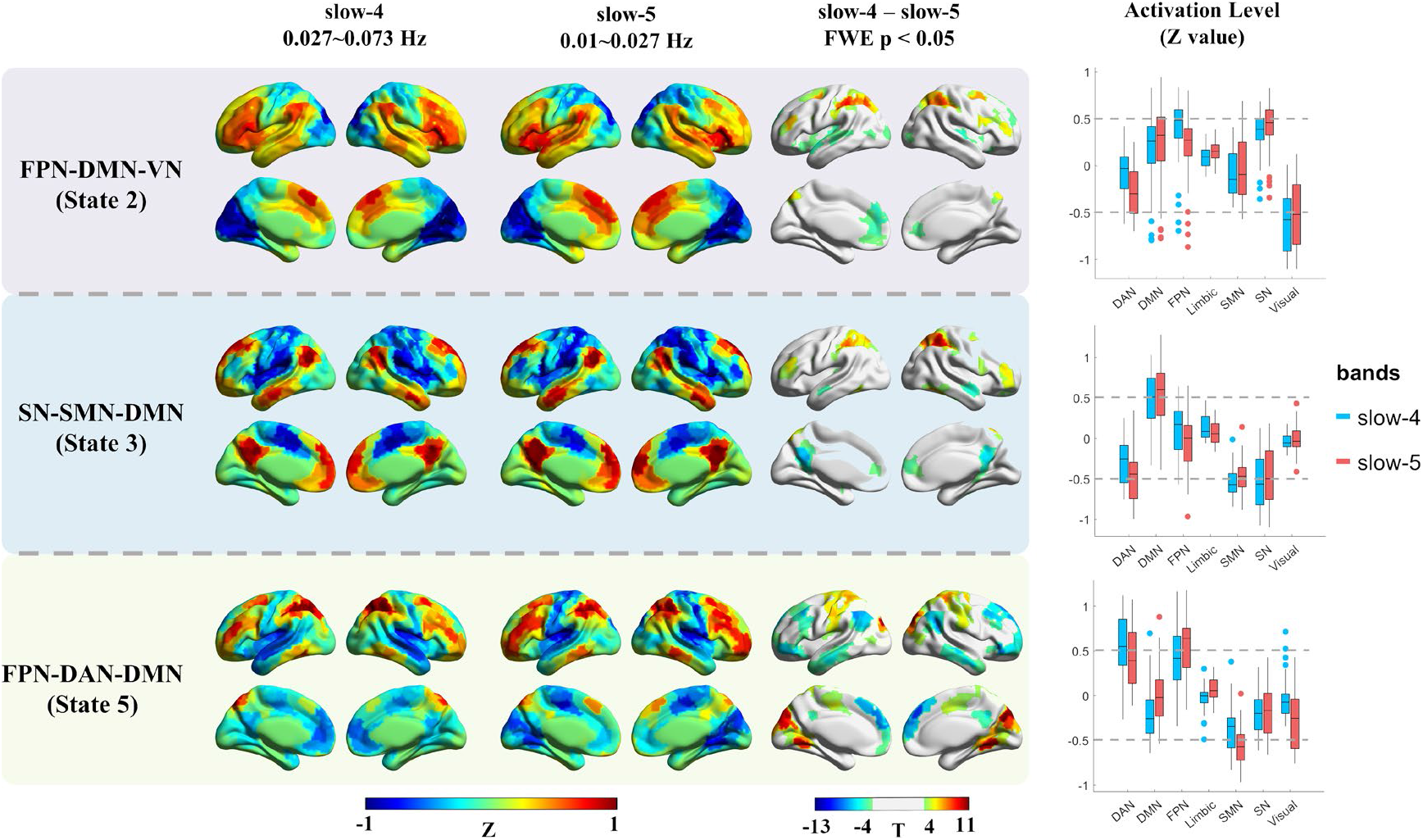
The frequency-specific effects between slow-4 and slow-5 within the HC group. The results of three states were presented, as the six CAP states were grouped into three pairs, and similar results were found within the pair. The first two columns show the cortical coactivations, and the color of each ROI indicates the activation deviation from its baseline level (Z-value). Paired t-test was performed for each state separately, and Bonferroni correction was used at the ROI level. The colorbar shows the T-value, and regions with P < 0.05 (FWE corrected) were presented in the third column. The last column shows the activation level of the seven networks in slow-4 and slow-5, and each point represents an ROI’s group averaged activation level from all 97 HC subjects. Abbreviations: DAN, dorsal attention network; DMN, default mode network; FPN, fronto-parietal network; SN, salience network; SMN, somatomotor network.

It can be observed that, State 1 and 2 were characterized by large activation deviation in the VN in both slow-4 and slow-5, then followed by SN and FPN. Compared with slow-5, slow-4 showed larger activation deviation in the bilateral middle frontal gyrus (FPN), and less activation deviation in the bilateral insula (SN) and dorsal attention network (DAN) in both State 1 and 2. Slow-4 also exhibited less anterior DMN activation in State 2. As for State 3 and 4, they were dominated by the SN, SMN and DMN in both slow-4 and slow-5, and slow-4 showed less activation deviation in the DMN and FPN. Besides, slow-5 was also dominated by the DAN, hence stronger activation deviation in the DAN was observed in slow-5. Finally, State 5 and 6 showed large DAN and FPN activation deviation in both slow-4 and slow-5, while slow-5 was also dominated by the SMN. In general, slow-5 showed an overall stronger activation deviation in the SMN, FPN and VN, and slow-4 showed a stronger activation deviation in the DMN.

In addition, frequency-specific activation differences have also been found in several subcortical regions after FDR correction (p < 0.005, FDR adjusted). Slow-4 exhibited an overall stronger subcortical activation deviation than slow-5 (Figure 5). Particularly, slow-4 showed stronger activations at the bilateral basal ganglia (caudate nucleus, putamen and globus pallidus) in State 6, stronger deactivations at the right caudate nucleus and globus pallidus in State 5, stronger activations at the bilateral amygdala in State 4, and stronger deactivations at the bilateral caudate nucleus, putamen and amygdala in State 3. Nevertheless, weaker deactivations at bilateral globus pallidus and left putamen were also found in slow-4 in State 1.

**Figure 5.**
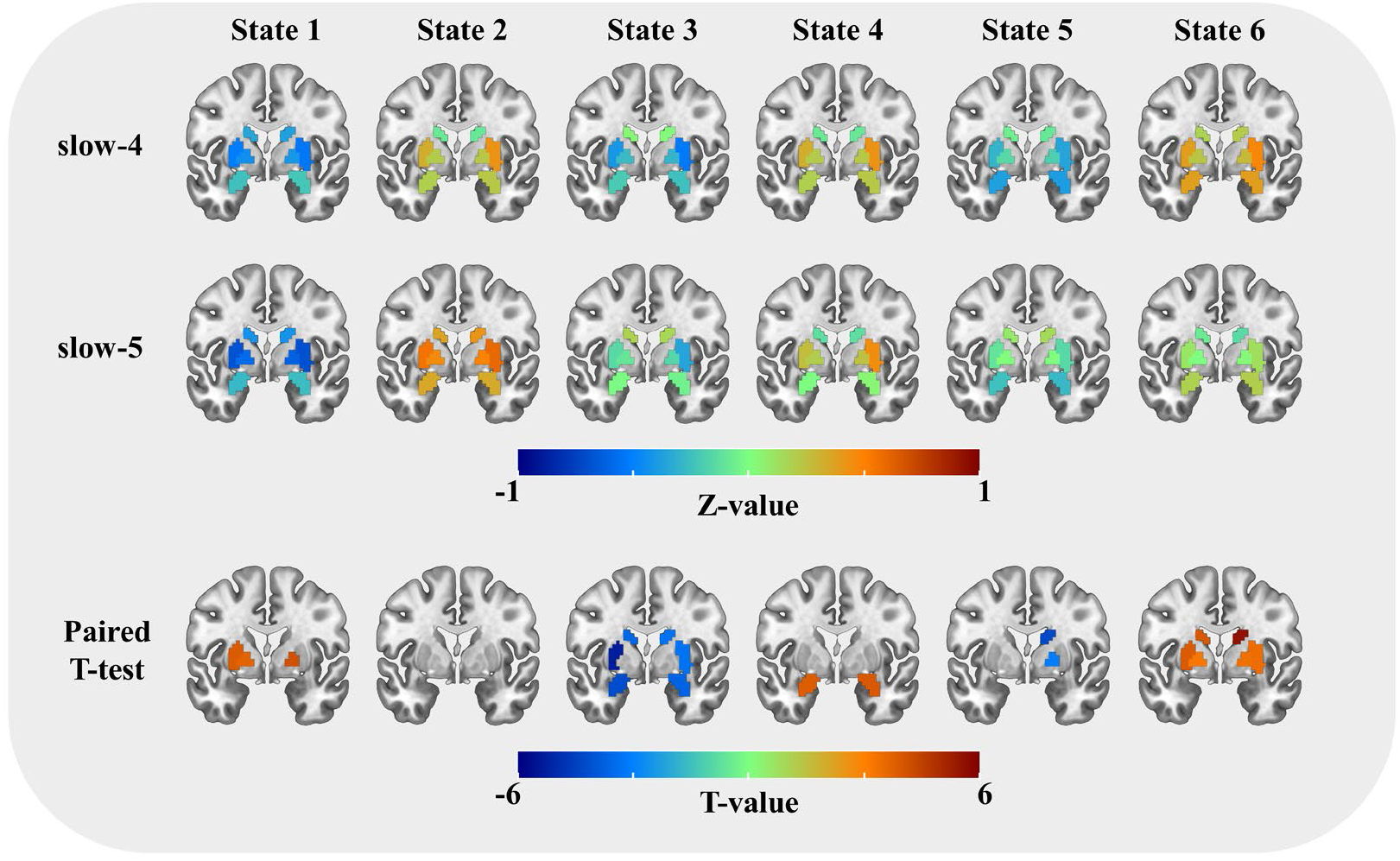
The subcortical activation differences between slow-4 and slow-5 within the HC group. The first two rows show the coactivations of the eight subcortical regions, and the color of each ROI indicates the activation deviation from its baseline level (Z-value). Paired t-test was performed for six states separately, and FDR correction was used at the ROI level. Regions with P < 0.005 (FDR adjusted) were presented in the last row, and the colorbar shows the T-value.

**Figure 6.**
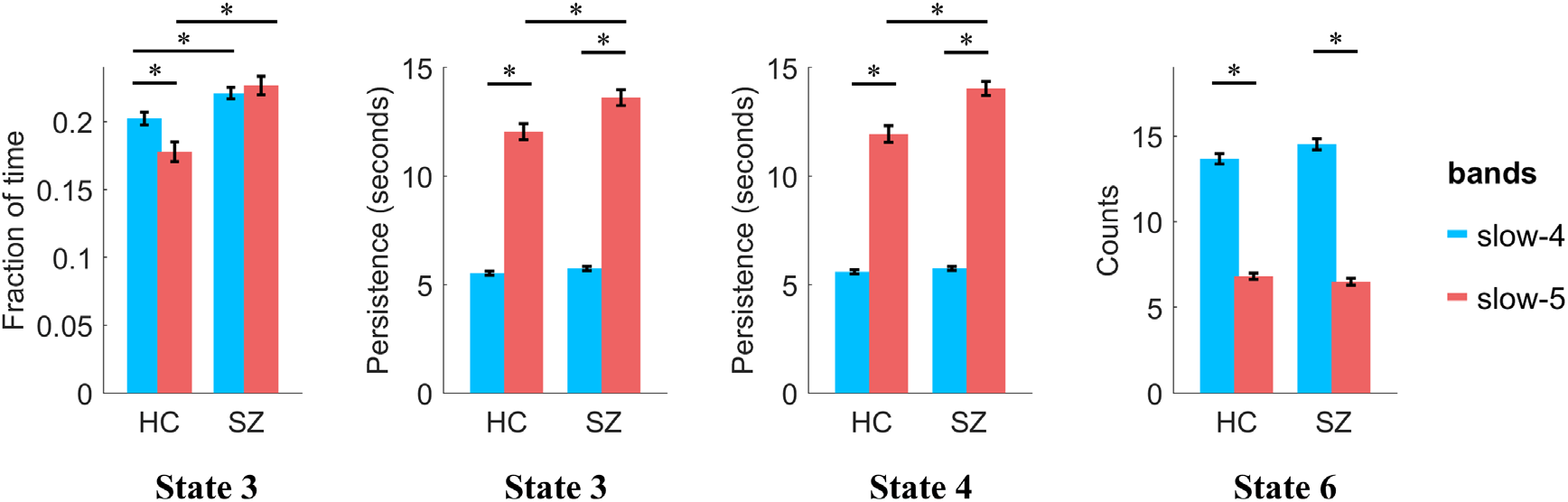
The post hoc results of repeated two-way ANOVA, only the results with significant interaction effects were compared. Between-group differences were compared using two-sample t-test, and between-frequency differences were compared using paired t-test. Age and gender were controlled for between-group comparisons, and FDR correction was performed to correct the multiple comparisons. Error-bar shows the standard error. * indicates p < 0.05 with FDR correction.

Next, the CAP temporal dynamics in slow-4 and slow-5 within the HC group were quantitatively compared using a paired t-test (supplementary Figure S7A). Compared with slow-5, all six states showed significantly shorter persistence and more counts in slow-4. More fraction of time in State 3 and State 4, and less fraction of time in State 5 and State 6 were observed in slow-4. In addition, the variation of fraction of time between the six states was also evaluated in slow-4 and slow-5 separately (supplementary Figure S7B). No between-state difference was found in slow-5, that each state occupied about 15% - 17% of the time. However, significant between-state differences were found in slow-4, e.g., State 3 and 4 showed more fraction of time (about 20 %) than the other four states, and State 6 accounted for the least amount of time.

### 3.3 Frequency-specific CAP dynamic alterations in SZ

The CAP differences between SZ and HC have been examined in the typical range before (Yang et al. 2021), then the frequency-specific alterations in slow-4 and slow-5 were further studied in this work. Two-way repeated-measures ANOVA was performed to estimate the main effects and interaction effects between group and frequency.

Significant group main effects were found in several states. SZ showed decreased fraction of time in State 1 and State 2, and increased fraction of time in State 3 and State 4. SZ also showed deceased persistence in State 2 and State 6, and increased persistence in State 3 and State 4. Finally, deceased counts in State 1 and State 2, and increased counts in State 3 and State 4 were observed in SZ. Significant frequency main effects on fraction of time were also found. Fraction of time increased in State 3 and State 4 and decreased in State 5 in slow-4. For persistence and counts, significant frequency main effects were obtained in all six states. As described before, higher-frequency (slow-4) CAPs showed shorter persistence and more counts than lower-frequency (slow-5). The detailed statistic results were presented in Supplementary Table S4.

Significant frequency-group interaction effects were found on fraction of time (P = 0.0014, F = 11.0473) in State 3, persistence in State 3 (P = 0.0084, F = 7.38) and State 4 (P = 4.48 × 10^−4^, F = 13.61), and counts in State 6 (P = 0.0465, F = 4.11). For post hoc results, we mainly reported the group differences. SZ showed increased fraction of time in State 3 in both slow-4 (P = 0.0058, T = 2.93) and slow-5 (P = 6.23 × 10^−6^, T = 4.90) than HC. Besides, SZ also showed increased persistence in both State 3 (P = 0.0052, T = 3.00) and State 4 (P = 1.04 × 10^−4^, T = 4.18) in slow-5 than HC. No group difference was obtained in either slow-4 or slow-5 on counts.

As for the classification results, the ROC (receiver operating characteristic) curves and their AUC (Area Under Curve) values were shown in Figure 7. Generally, the spatial features resulted in a higher classification accuracy than the temporal features, and combining slow-5 and slow-4 would increase the AUC than only using the typical range. The best classification results were obtained by combining the spatial features from both slow-5 and slow-4, with AUC = 0.9630, Accuracy = 0.8913, Sensitivity = 0.8986 and Specificity = 0.8841. The detailed results can be found in Supplementary Table S5 and Table S6.

**Figure 7.**
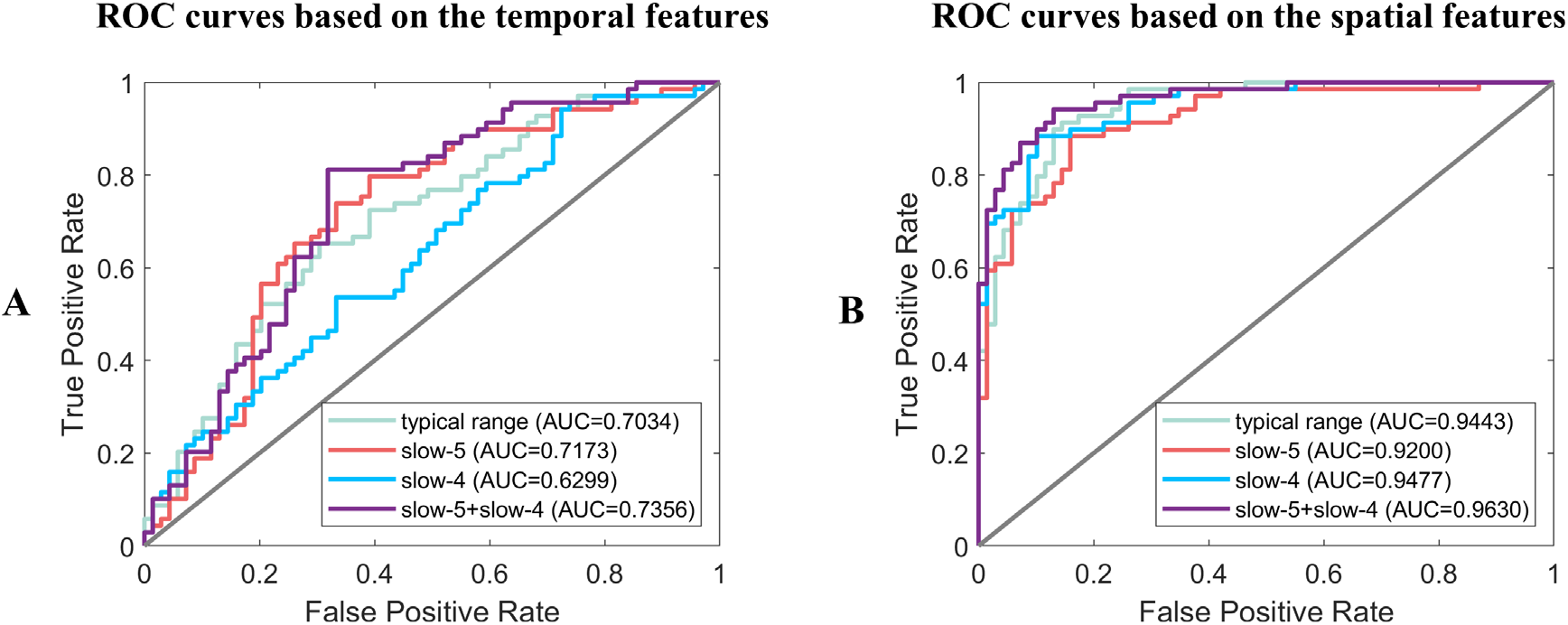
The ROC curves and AUC values of classification results based on the (A) temporal features and (B) spatial features in the typical range, slow-5, slow-4 separately and combined slow-5 with slow-4.

## 4. Discussion

This work systematically investigated the frequency-specific coactivation patterns across slow-2 to slow-5 and compared their spatial and temporal characteristics with the results obtained from the typical range. Particularly, slow-4 and slow-5 were further compared within healthy subjects, and the frequency-specific CAP dynamic alterations in schizophrenia patients were studied. Generally, both the high-order and primary networks can be observed across slow-5 to slow-3 except slow-2, and the CAP state persisted shorter and occurred more frequently at a higher frequency band. In addition, stronger subcortical coactivations and co-deactivations were observed in slow-4, while large-scale function networks such as DAN showed stronger coactivations and co-deactivations in slow-5. Furthermore, schizophrenia patients showed frequency-specific alterations in slow-4 and slow-5, and combining slow-4 and slow-5 increased the classification accuracy.

### 4.1 Spatial and temporal properties of CAPs at different frequency bands

The coactivation patterns in the typical range (0.01 - 0.08 Hz) have been demonstrated in our previous study (Yang et al. 2021), and the four sub-frequency bands were further studied in this study. Compared with the typical range, slow-5 (0.01 - 0.027 Hz), slow-4 (0.027 - 0.073 Hz) and slow-3 (0.073 - 0.198 Hz) showed similar spatial patterns, which were characterized by typical functional networks. Consistent with previous studies (Huang et al. 2020; Zhang et al. 2020), opposite CAP pairs were also found in the sub-frequency band, suggesting the antagonistic relationships between these intrinsic networks widely exist at different sub-frequency bands. However, the CAP states in slow-2 (0.198 - 0.25 Hz) were unlike that of the typical range, brain regions belonging to the same network were not coactivated together or mixed with other networks. A previous study has found that slow-2 mainly oscillates within white matter rather than grey matter (Zuo et al. 2010). Gohel and Biswal (Gohel and Biswal 2015) evaluated the seed-based correlation maps from slow-5 to slow-1, and they found the spatial extent of slow-2 was significantly reduced compared with slow-5/4/3. Together, these findings indicate the attenuated intrinsic functional associations in slow-2. The reason might be that the resting-state brain and intrinsic functional networks were mainly activated at the low-frequency, and there were also more physiological noises at the higher frequency (Cordes et al. 2001; Chen and Glover 2015).

As for the CAP dynamics, persistence and counts changed monotonically with the increased frequency band. Particularly, persistence decreased, and counts increased for all the six states from slow-5 to slow-2, suggesting that the higher frequency led to unstable state maintenance. First, the higher frequency could cause more frequent BOLD fluctuations, hence the volume-to-volume state maintenance would decrease, and the between-state transition would increase. As the fraction of time was similar across different frequency bands, shorter persistence would lead to more counts. Besides, the higher frequency BOLD signal involved more noises (Cordes et al. 2001; Chen and Glover 2015), which might affect the coactivation profile and result in more between-state transitions, and shorten the persistence.

### 4.2 Specific spatial and temporal configurations of CAPs in slow-4 and slow-5

Previous studies have found that the neural oscillations of the human brain are frequency-dependent. Particularly, large-scale functional networks (e.g., DMN and FPN) integrate remote brain regions with long-distance interactions (Salvador et al. 2005), and these functional processes are primarily achieved by a lower-frequency band (Buzsaki and Draguhn 2004; Penttonen, Buzsáki, and Systems 2003). On the contrary, subcortical regions are spatially compact and dominated by local neural activities (Salvador et al. 2005), and these short-range connections work in a higher-frequency band (Buzsaki and Draguhn 2004).

In our results, generally the whole-brain spatial patterns of the six CAP states in slow-4 and slow-5 were similar to the typical range, which were characterized by the coactivation or co-deactivation of large-scale intrinsic networks, while frequency-specific coactivation profiles were still found between slow-4 and slow-5. Stronger DAN, DMN and FPN activation deviations were observed in slow-5 in several CAP-states. Especially, DAN showed larger activation deviation in slow-5 across State 1 to State 4, suggesting the activity of DAN mainly fluctuates at the lower frequency band. Slow-5 showed stronger DMN activation deviation in State 3 and 4, and stronger anterior DMN activation in State 2. Previous studies have also found greater ALFF/fALFF in several DMN regions in slow-5 (Han et al. 2011; Wang, Kong, et al. 2016). Subcortical regions showed larger activation deviations in slow-4 for most CAP states, for instance, slow-4 showed stronger activation at bilateral basal ganglia in State 6, which was consistent with previous findings that stronger basal ganglia ALFF/fALFF in slow-4 (Zuo et al. 2010). The above results support the previous findings that large-scale functional networks are mainly mediated by lower-frequency connections, while higher-frequency activities are linked with subcortical systems (Buzsaki and Draguhn 2004; Han et al. 2011; Wang, Kong, et al. 2016).

However, different compared with previous results, stronger FPN activation deviation has been observed in slow-4 in State 1 and 2, and stronger DMN activation deviation has been found in slow-4 in State 5 and 6. Furthermore, a few subcortical regions (globus pallidus and putamen) also showed weaker activation deviations in slow-4 in State 1. It is worth noting that, previous conclusions were mainly drawn from static studies, while the frequency-specific properties might change dynamically, and the subcortical-related high-frequency band (0.3032–0.4545 Hz) studied before is (Salvador et al. 2005) even higher than slow-2 (0.198 - 0.25 Hz). While both slow-5 (0.01 - 0.027 Hz) and slow-4 (0.027 - 0.073 Hz) still belong to the typical low-frequency range (0.01 - 0.08 Hz), which might be the reason why the subcortical and large-scale networks were not always stronger in slow-4 or slow-4 across all states. Therefore, these results suggest that within the typical low-frequency range, although both slow-4 and slow-5 showed similar coactivation patterns, the large-scale networks and subcortical regions might still be mediated by different frequency bands at specific periods and brain states.

As for the temporal domain, slow-4 showed significantly shorter persistence and more counts across all the six CAPs. Due to the long persistence and high within-state transition probability, fewer between-state transitions occurred in slow-5, suggesting the integration of large-scale networks requires sufficient time to maintain a relatively stable state and execute specific functions based on the lower-frequency signals. Besides, an unbalanced between-state fraction of time was found in slow-4 but not in slow-5 (Figure S7B). State 3 and 4 showed more fraction of time than the other four CAPs in slow-4, and State 5 and 6 showed the least fraction of time. Hence, slow-4 showed a significantly increased fraction of time in State 3 and 4, and decreased fraction of time in State 5 and 6. The unbalanced between-state time allocation and more frequent between-state transitions together suggest the richer temporal dynamics in slow-4.

### 4.3 Frequency-specific CAP differences between SZ and HC in slow-4 and slow-5

Previous studies have shown that SZ patients are not only characterized by frequency-specific changes (Gohel et al. 2018; Yu et al. 2013; Zhang, Yang, and Cai 2020) or temporal dynamic changes (Du et al. 2017; Kottaram et al. 2018), but also frequency-specific dynamic alterations (Zou and Yang 2019; Luo, He, et al. 2020). Our previous study has demonstrated the altered CAP dynamics between SZ and HC in the typical range (Yang et al. 2021), and the frequency-specific alterations in SZ were further studied in the current work. In general, a similar trend of case-control differences across the six states was observed at both slow-4, slow-5 and the typical range. Consistent with the typical range, SZ patients showed less fraction of time in the FPN-DMN state (State 1 and 2) and more in the SN-DMN state (State 3 and 4) in both slow-4 and slow-5. The FPN, DMN and SN have been widely reported abnormal in psychiatric disorders as triple-network model (Manoliu et al. 2014; Menon 2011; Supekar et al. 2019), and our results further showed the altered triple-network dynamics in SZ patients exist in both slow-4 and slow-5. Besides, frequency-specific alterations were also found between slow-4 and slow-5. Particularly significant group-frequency interactions in the SN-DMN state were obtained on all the three CAP dynamics (fraction of time, persistence and counts), and increased persistence in the SN-DMN state was only obtained in slow-5 but not in slow-4 nor the typical range. The reason might be that slow-5 was characterized by stronger DMN activation deviation in the SN-DMN state, and the more active spatial foundation provides richer temporal dynamics, enabling the discovery of more distinguished disease alterations. Furthermore, combining the features from both slow-4 and slow-5 have increased the diagnose accuracy of schizophrenia patients than only use the typical range. Previous studies have also shown the increased classification accuracy by combing slow-4 and slow-5 (Huang et al. 2019; Tian et al. 2020). Together, these results suggest that frequency-dependent dynamic information contains in multi-frequency bands, and could help to identify frequency-specific disease alterations.

### 4.4 Limitations

In this study, we used frequency divisions from slow-5 to slow-2 as was described by Buzsaki (Buzsaki and Draguhn 2004) and used by Zuo and colleagues (Zuo et al. 2010) in fMRI. Although, this method has been widely adapted in fMRI that uses unequal ranges of frequency bandwidths, recent studies have shown a wavelet-based method with higher sensitivity and reproducibility to obtain the sub-frequency bands (Luo, Wang, et al. 2020). Future studies should validate and compare the frequency-dependent CAPs obtained by wavelet transform and fast Fourier transform. Besides, the effects of physiological noises such as head movement, cardiac and respiratory motion were not fully studied. Although, subjects with large headmotion were excluded before the analysis, and the 24 headmotion parameters (Friston et al. 1996) were regressed from the BOLD signal, headmotion could still affect the dynamic functional connectivity (Nalci, Rao, and Liu 2019; Laumann et al. 2017). Whether and how the headmotion would influence the spatial and temporal characteristics of CAPs remains further study. The cardiac and respiratory motions were not recorded during the scan, and have not been corrected from the time series using methods such as RETROICOR (Glover, Li, and Ress 2000). While slow-3 and slow-2 might be involved with these high-frequency noises, hence their effects on CAPs cannot be studied systematically in current work.

## 5. Conclusions

This study proved that the resting-state CAP states showed frequency-specific spatial and temporal characteristics. In summary, from slow-5 to slow-2, the spatial patterns varied from intrinsic functional networks to irregular configurations, and the CAP state became more unstable and frequently changed when the frequency band increased, which caused shorter persistence and more counts. Besides, our results supported that, the large-scale network integration relies more on lower-frequency oscillations (slow-5) and the subcortical regions activate more in a relative higher-frequency band (slow-4), from a dynamic point of view. Finally, frequency-dependent dynamic changes in schizophrenia patients were also observed between slow-5 and slow-4. Our results could provide more information about the functional dynamic brain, and help to understand the frequency-specific pathological mechanisms of psychiatric disorders.

## Data and code availability statements

The data used in this study is not publicly available due to privacy or ethical restrictions. The code that supports the findings of this study will be made available upon request from the corresponding author.

## Acknowledgements

We are grateful to all the patients and volunteers of this study as well as the staffs at the Wuxi Mental Health Center for their help with participant recruitment and data collection. This work was supported by the National Natural Science Foundation of China (NSFC) grant (No. 62071109 to C.M., No. 81871081 and 81301148 to L.T., No. 61871420 to B.B.,).

