## supplementary materials for "Frequency-specific coactivation patterns in resting-state and their alterations in schizophrenia: an fMRI study"

**^*^ Correspondence:**

**Methods and Results**

**Classification Analysis**

LIBSVM (Chang & Lin, 2011) was used to train the SVM classifier, with a linear kernel and C = 1. Eight classification models were built in this study, the temporal and spatial characteristics of the six CAP states were fed into the SVM as features independently for the typical range, slow-5, slow-4 and slow-5 + slow-4. In detail, each subject has 18 (3*6) temporal features which include fraction of time, persistence and counts of the six CAPs, and 2448 (408*6) spatial features which represent the 408 ROIs’ activation level of the six CAPs.

A nested-LOOCV (leave-one-out cross-validation) SVM-Fscore method (Chen & Lin, 2006) was used to get the model with the highest accuracy, and F-score was used to select features. The feature number was tested from 20 to 1500 with a step length of 20 for the spatial features, and from 1 to 18 (36 for slow-5 + slow-4) with a step length of 1 for the temporal features. For each LOOCV iteration, the F-score of all features was calculated and ranked within the training set, where a larger F-score indicates larger group differences. Then, the smallest step that achieved the highest accuracy was chosen and the corresponding classification results were reported. Accuracy, sensitivity, specificity, and area under curve (AUC) were calculated to measure the performance of the classifier.

**Statistics**

Instead of using two-way repeated ANOVA to estimate the group-frequency interactions and group main effects, simple effect analyses have also been performed for all the six CAP states. The temporal CAP differences between SZ and HC were compared in slow-5 and slow-4 separately, using a two-sample t-test with age and gender as covariates, and FDR correction was performed to account for the multiple comparisons.

As shown in Figure S10, SZ showed a consistently decreased fraction of time in the FPN-DMN-VN state (State 1 and 2), and an increased fraction of time in the SN-SMN-DMN state (State 3 and 4) in both slow-4 and slow-5. Moreover, SZ showed shorter persistence in the FPN-DMN-VN state (State 1 and 2) and FPN-DAN-DMN state (State 6) in slow4, while longer persistence in the SN-SMN-DMN state (State 3 and 4) was found in slow-5. As for the counts, SZ showed more counts in State 4 in slow-4, fewer counts in State 1 and 2 and more counts in State 3 in slow-5. No significant difference was found for transition probability in either slow-4 or slow-5 after FDR correction.

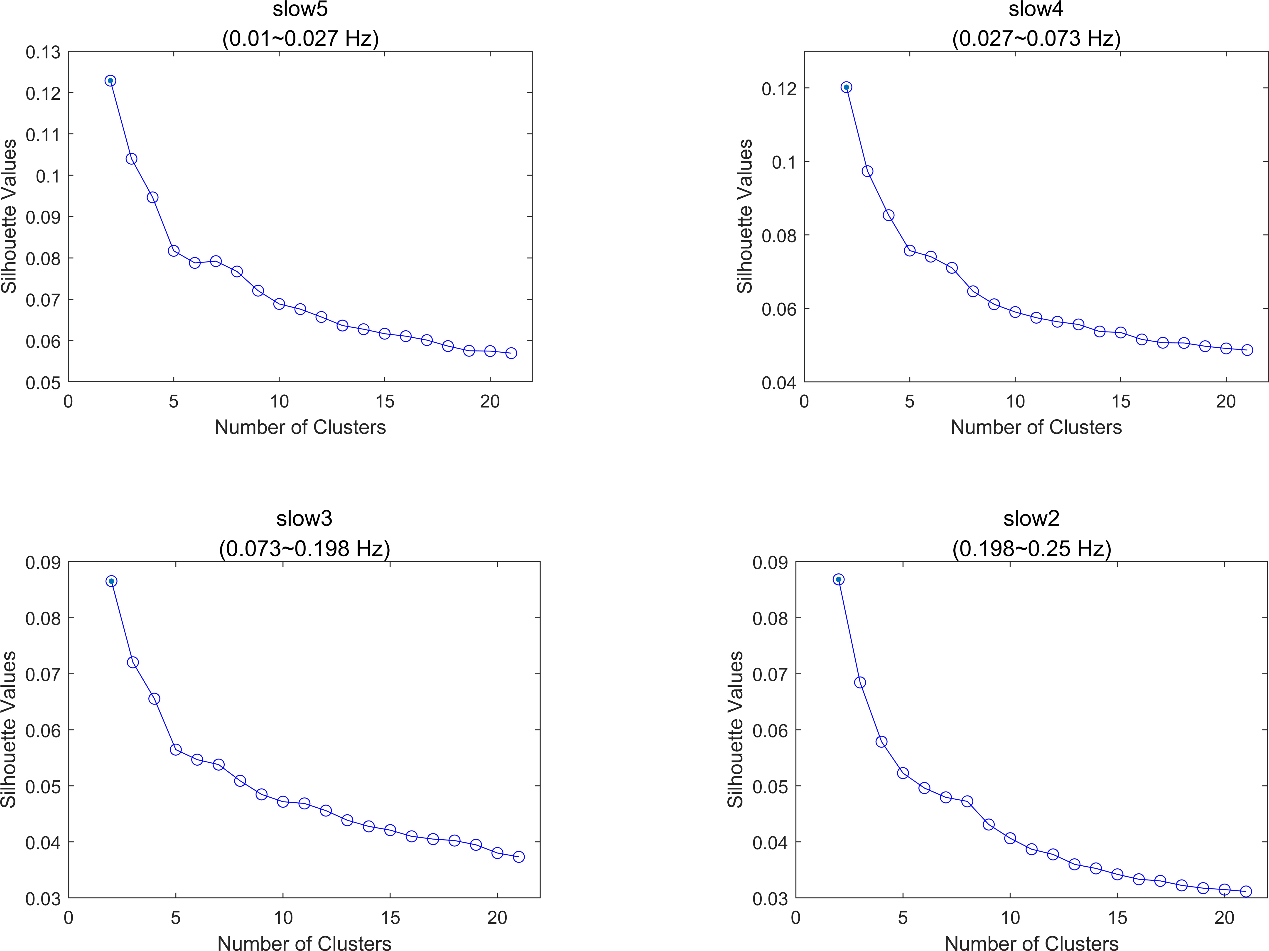

**Figure S1.** The clustering curves of the four sub-frequency bands. Silhouette value was calculated from k = 2 to k = 21 with step length = 1, and the silhouette values were monotonically decreasing with the increase of k. The elbow point for the four curves was around 5 to 7, and k = 6 was chosen in the manuscript.

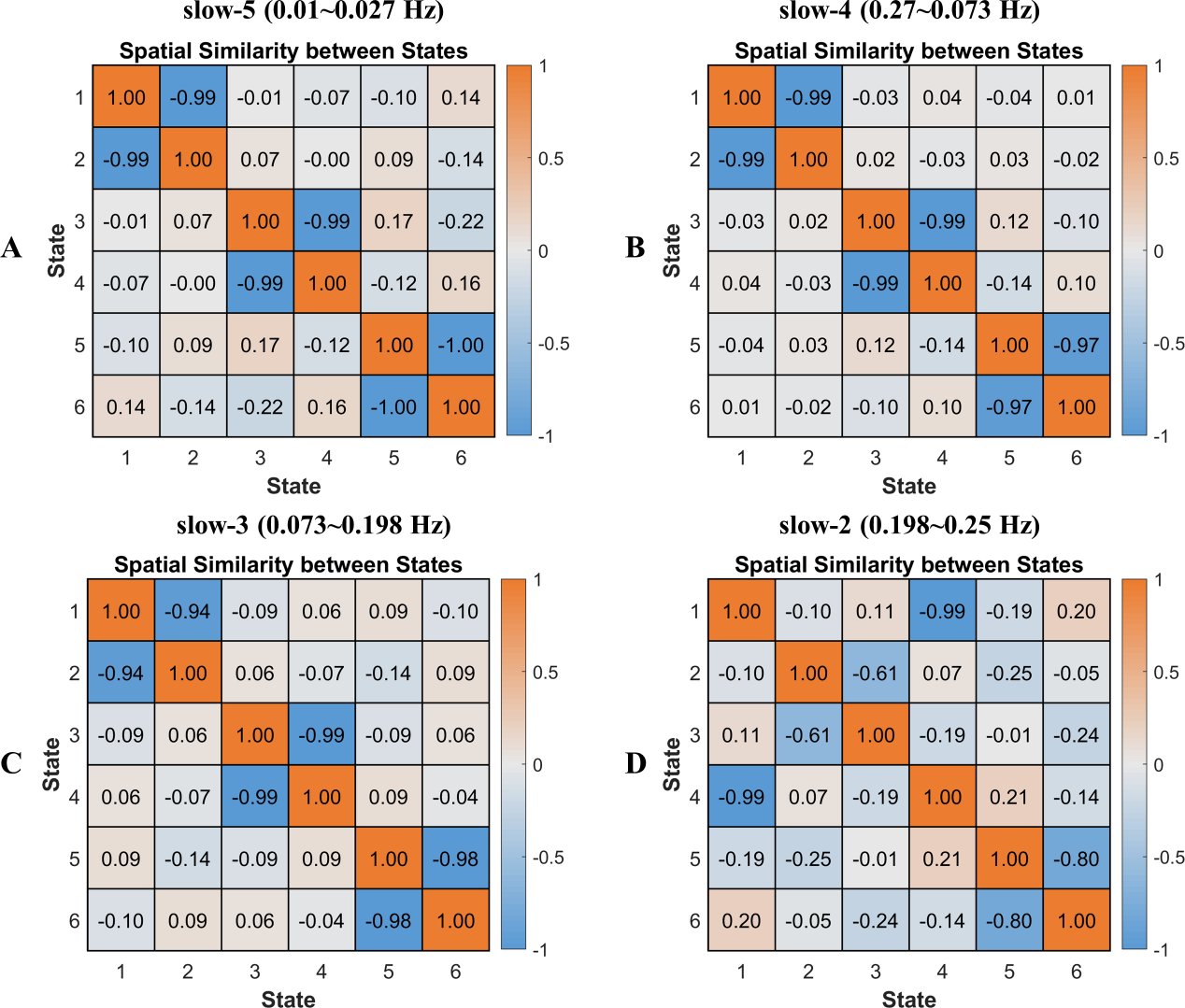

**Figure S2.** The CAP spatial similarity within (A) slow-5, (B) slow-4, (C) slow-3 and (D) slow-2. Pearson correlation was calculated to measure their spatial similarity. The colorbar shows the R-value.

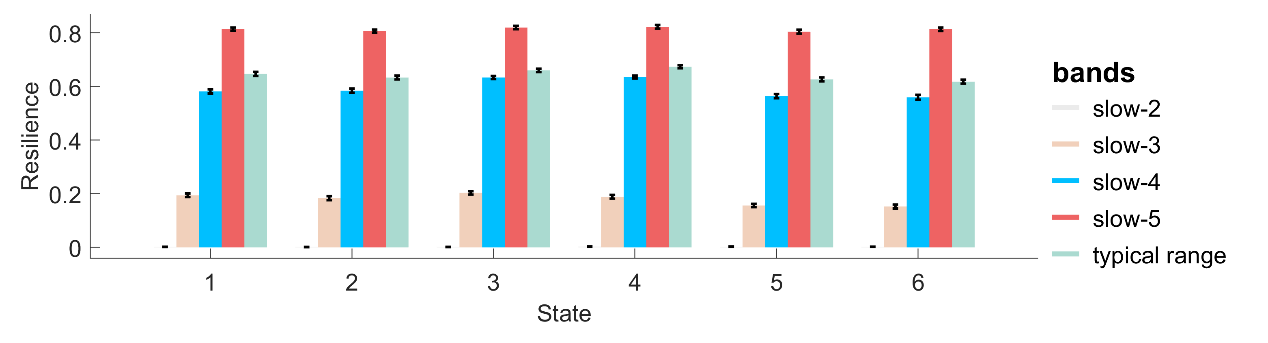

**Figure S3.** The resilience (within-state transition probability) of HC across slow-5 to slow-2.

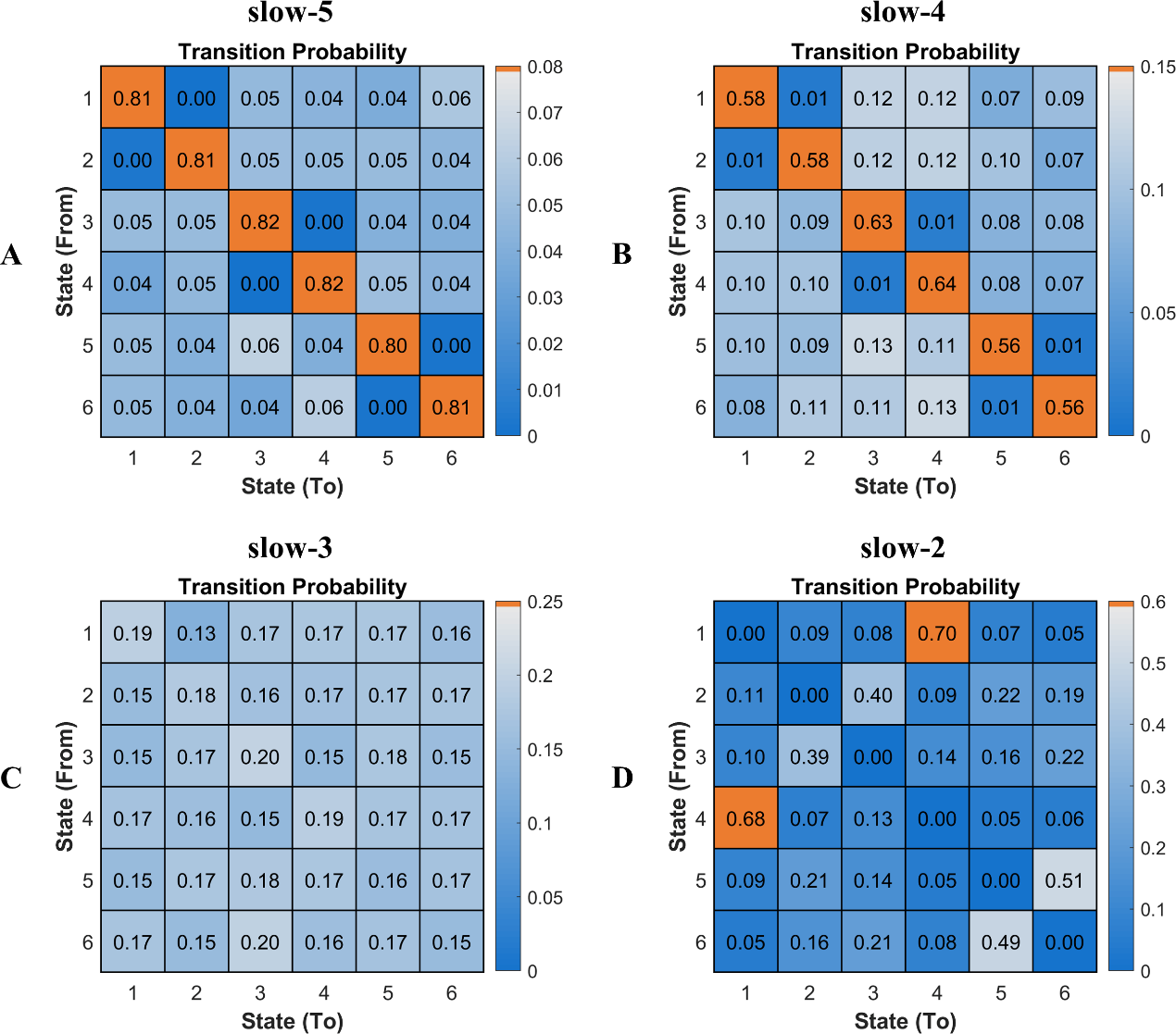

**Figure S4.** The transition probability matrix of HC across slow-5 to slow-2.

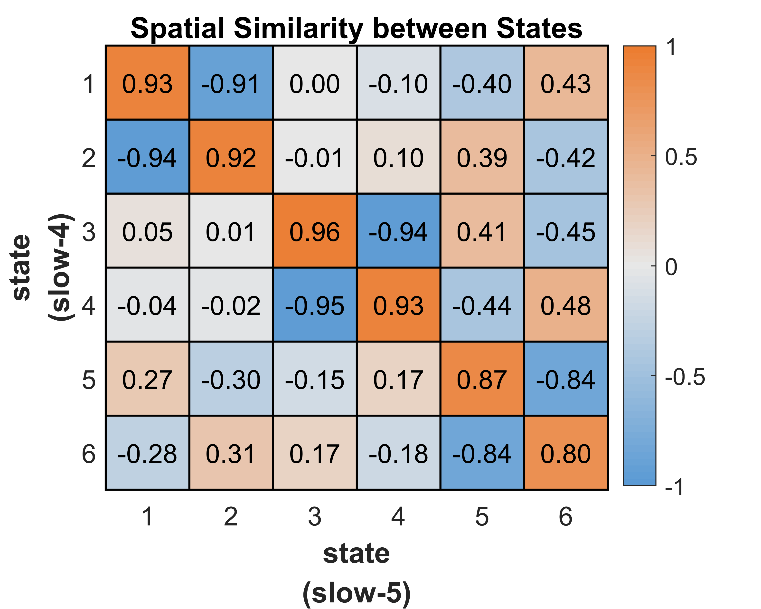

**Figure S5.** The CAP spatial similarity between slow-4 and slow-5. Pearson correlation was calculated to measure their spatial similarity. The colorbar shows the R-value.

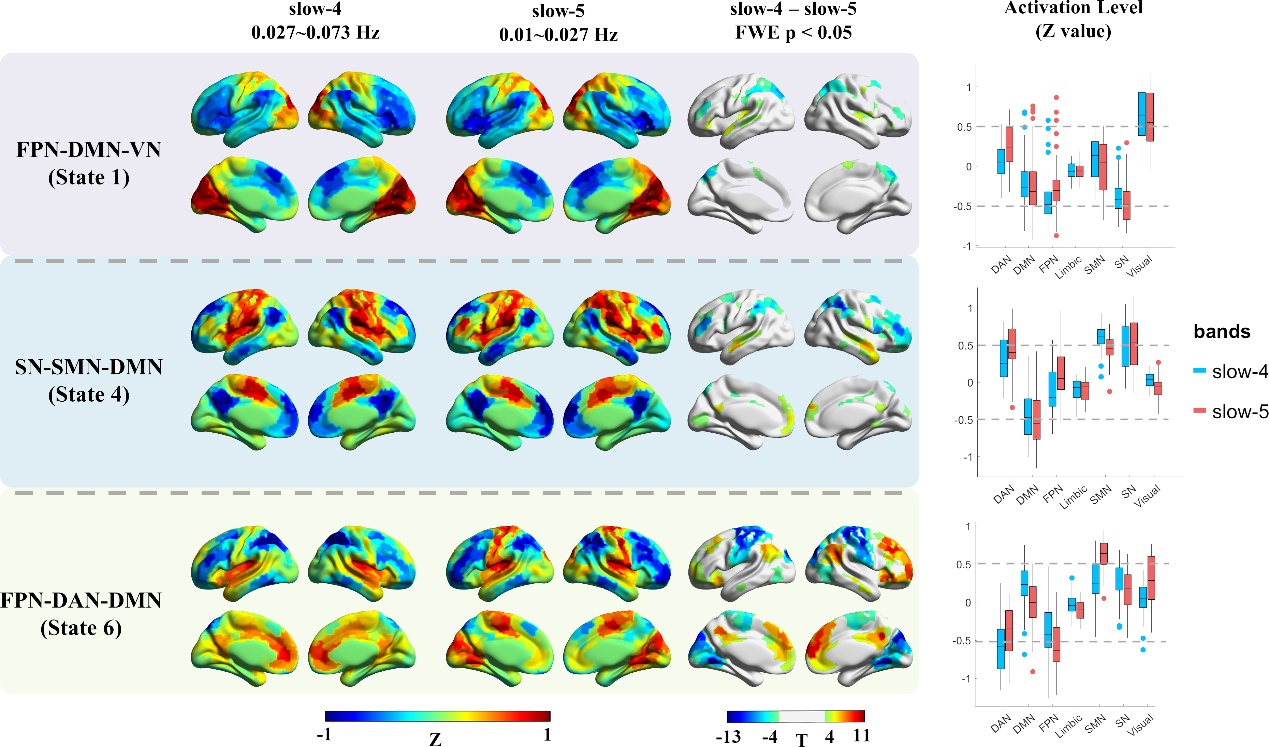

**Figure S6.** The frequency-specific effects between slow-4 and slow-5 within the HC group. The results of three states were presented, as the six CAP states were grouped into three pairs, and similar results were found within the pair. The first two columns show the cortical coactivations, and the color of each ROI indicates the activation deviation from its baseline level (Z-value). Paired t-test was performed for each state separately, and Bonferroni correction was used at the ROI level. The colorbar shows the T-value, and regions with P < 0.05 (FWE corrected) were presented in the third column. The last column shows the activation level of the seven networks in slow-4 and slow-5, and each point represents an ROI’s group averaged activation level from all 97 HC subjects.

**Abbreviations:** DAN, dorsal attention network; DMN, default mode network; FPN, fronto-parietal network; SN, salience network; SMN, somatomotor network.

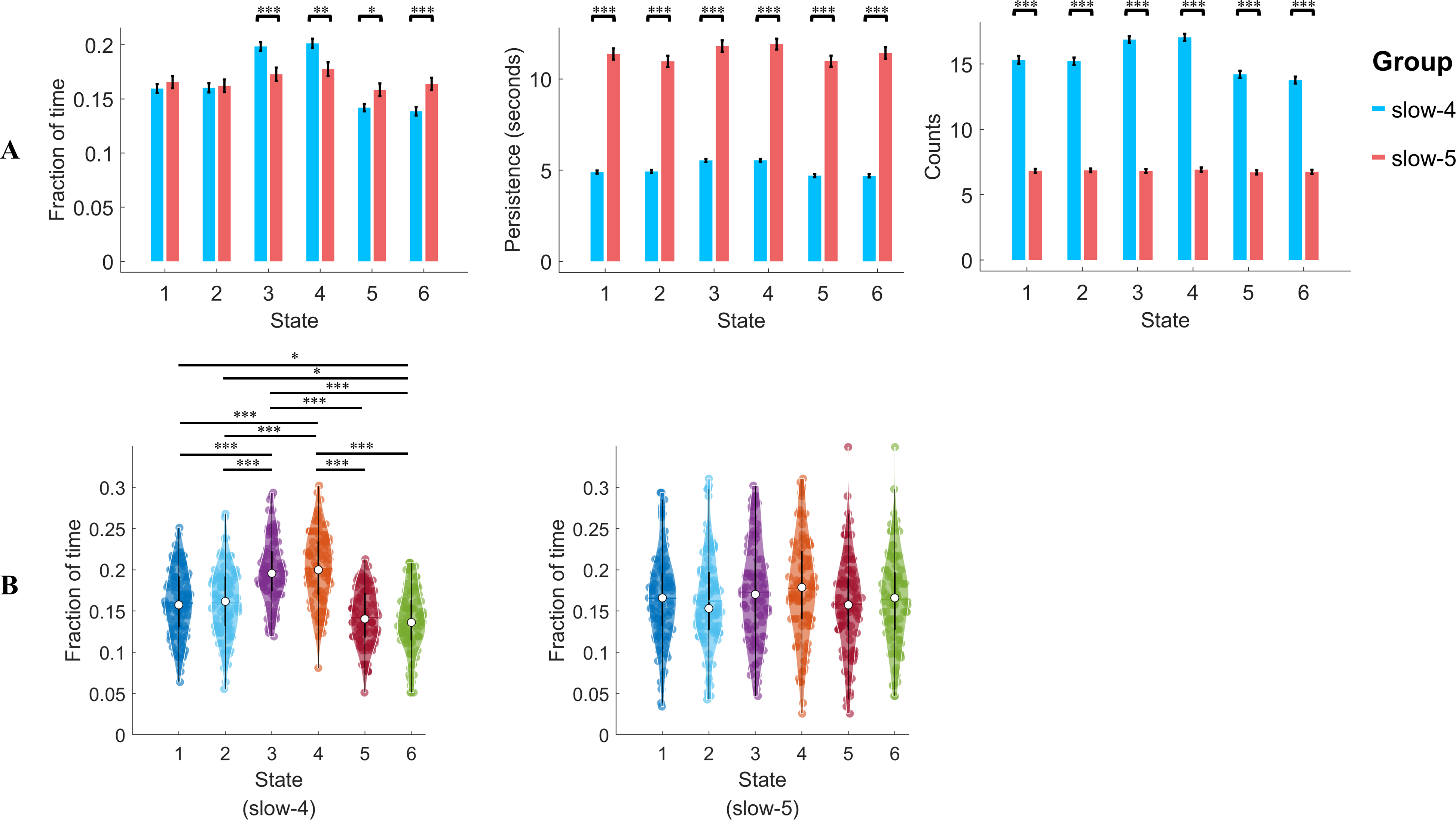

**Figure S7.** (A) The state temporal dominances (fraction of time, persistence, counts) under slow-4 and slow-5 within the HC group, and compared using paired t-test. (B) The fraction of time differed between six states at slow-4 but not at slow-5. Paired t-test was performed between each pair of states separately. Error-bar shows the standard error. * indicates p < 0.05, and ** indicates p < 0.005, and *** indicates p < 0.0005 separately, with FDR correction.

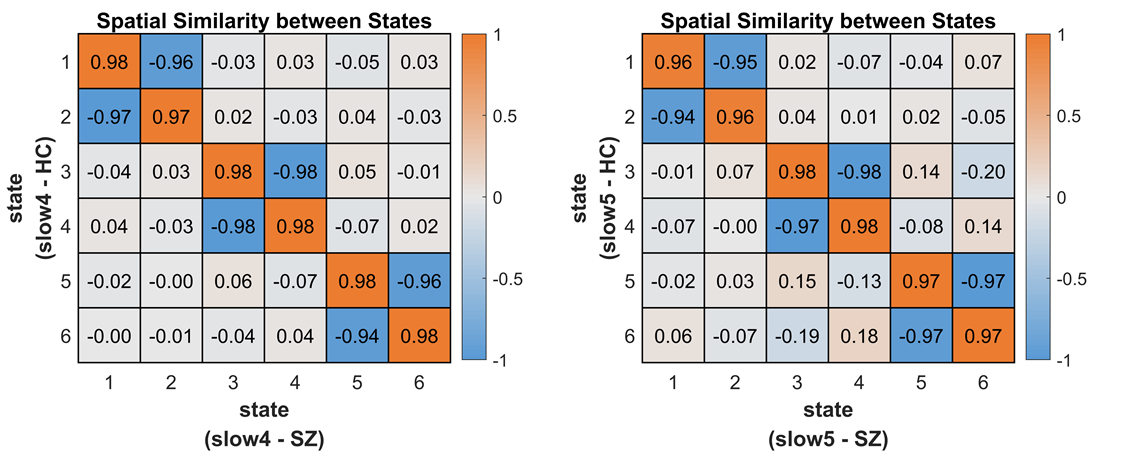

**Figure S8.** The CAP spatial similarity between SZ and HC in slow-4 and slow-5. Pearson correlation was calculated to measure their spatial similarity. The colorbar shows the R-value.

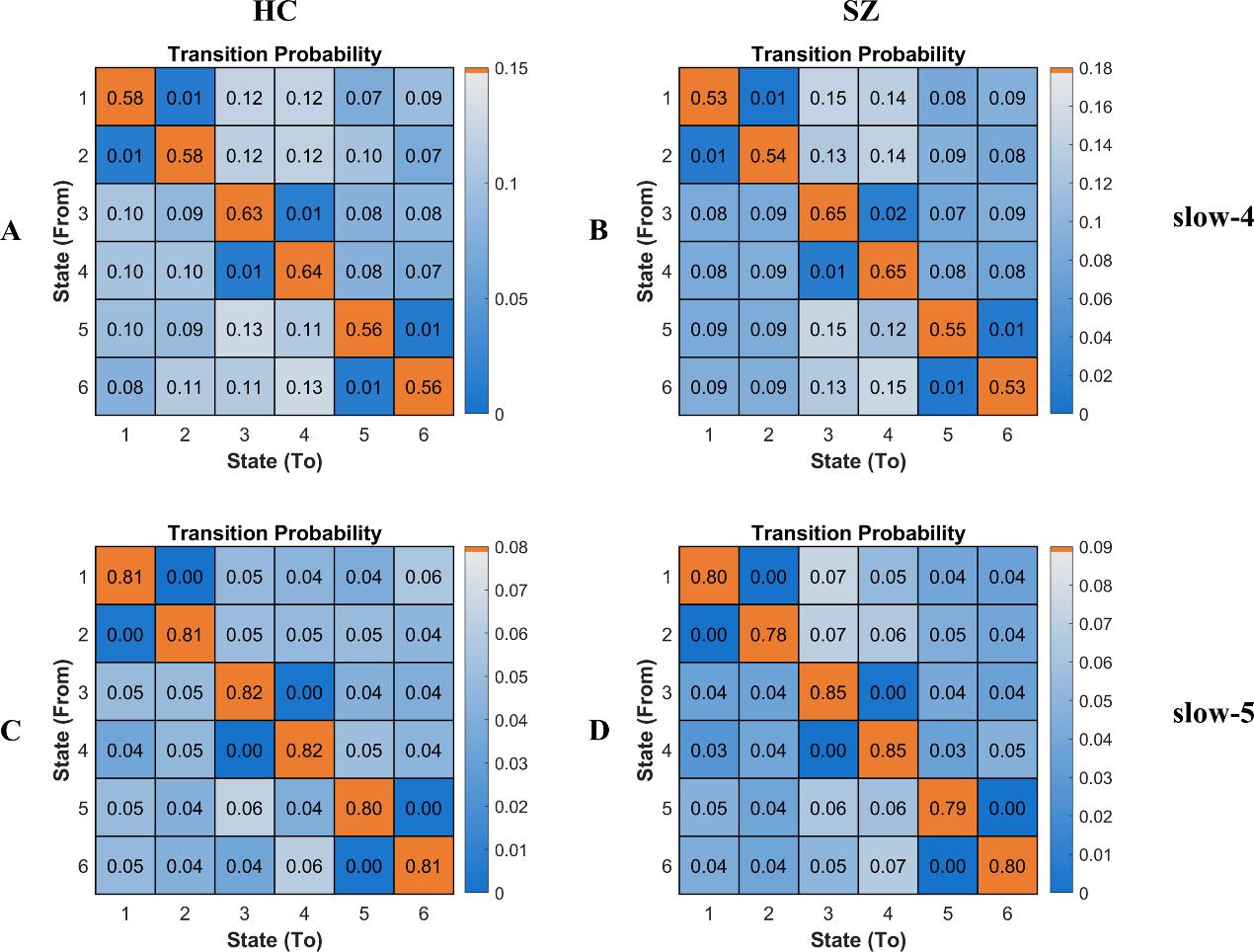

**Figure S9.** The transition probability matrix of HC and SZ in slow-4 and slow-5.

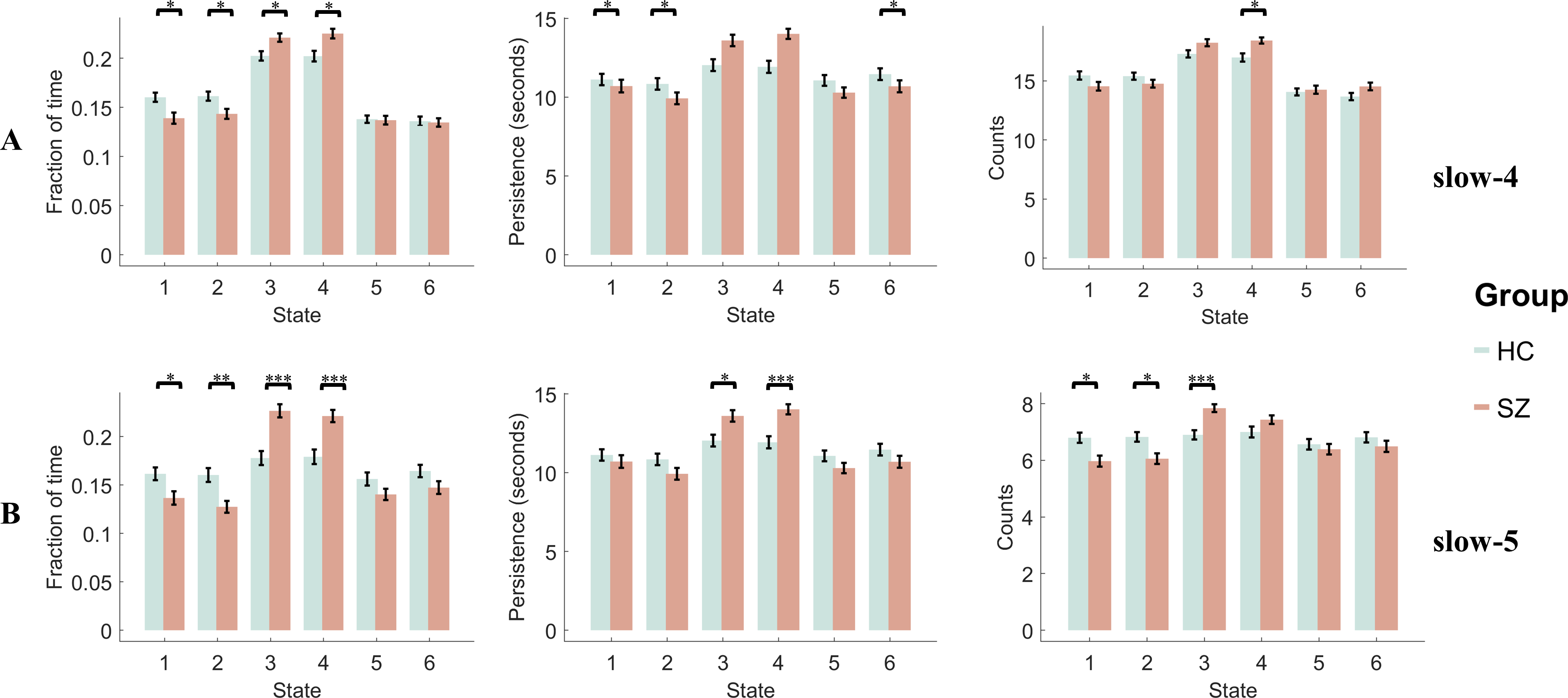

**Figure S10.** The state temporal dominances (fraction of time, persistence, and counts) under (A) slow-4 and (B) slow-5, and compared between SZ and HC using a two-sample t-test. Error-bar shows the standard error. * indicates p < 0.05, and ** indicates p < 0.005, and *** indicates p < 0.0005 separately, with FDR correction.

**Table S1.** The paired t-test results for CAP dynamic differences between slow-4 and slow-5 in the HC group.

| **Fraction of time** | **T value** | **P value (FDR adjusted)** |
| --- | --- | --- |
| State 1 | -1.0788 | 0.3401 |
| State 2 | -0.3402 | 0.7345 |
| State 3 | 4.6502 | 0.0001^*^ |
| State 4 | 3.6518 | 0.0008^*^ |
| State 5 | -2.6540 | 0.0140^*^ |
| State 6 | -4.0468 | 0.0003^*^ |
| **Persistence** | **T value** | **P value (FDR adjusted)** |
| State 1 | -21.4037 | <0.0001^*^ |
| State 2 | -20.6626 | <0.0001^*^ |
| State 3 | -20.7937 | <0.0001^*^ |
| State 4 | -21.8191 | <0.0001^*^ |
| State 5 | -21.0289 | <0.0001^*^ |
| State 6 | -21.1770 | <0.0001^*^ |
| **Counts** | **T value** | **P value (FDR adjusted)** |
| State 1 | 28.5528 | <0.0001^*^ |
| State 2 | 30.6306 | <0.0001^*^ |
| State 3 | 41.5672 | <0.0001^*^ |
| State 4 | 32.7739 | <0.0001^*^ |
| State 5 | 26.2321 | <0.0001^*^ |
| State 6 | 22.1921 | <0.0001^*^ |

^*^ indicates the FDR-adjusted p-value < 0.05.

**Table S2.** The two-sample t-test results for CAP dynamics differences between SZ and HC in slow-5.

| **Fraction of time** | **T value** | **P value (FDR adjusted)** |
| --- | --- | --- |
| State 1 | -2.6171 | 0.0148^*^ |
| State 2 | -3.4805 | 0.0014^*^ |
| State 3 | 4.8993 | 1.6367e-05^*^ |
| State 4 | 4.2655 | 0.0001^*^ |
| State 5 | -1.7698 | 0.0790 |
| State 6 | -1.8570 | 0.0786 |
| **Persistence** | **T value** | **P value (FDR adjusted)** |
| State 1 | -0.7672 | 0.4443 |
| State 2 | -1.7484 | 0.1550 |
| State 3 | 2.9974 | 0.0097^*^ |
| State 4 | 4.1817 | 0.0003^*^ |
| State 5 | -1.6401 | 0.1550 |
| State 6 | -1.4422 | 0.1819 |
| **Counts** | **T value** | **P value (FDR adjusted)** |
| State 1 | -3.0981 | 0.0059^*^ |
| State 2 | -3.0284 | 0.0059^*^ |
| State 3 | 4.4068 | 0.0001^*^ |
| State 4 | 1.7656 | 0.1196 |
| State 5 | -0.6590 | 0.5110 |
| State 6 | 1.1828 | 0.2568 |

^*^ indicates the FDR-adjusted p-value < 0.05.

**Table S3.** The two-sample t-test results for CAP dynamic differences between SZ and HC in slow-4.

| **Fraction of time** | **T value** | **P value (FDR adjusted)** |
| --- | --- | --- |
| State 1 | -2.9029 | 0.0086^*^ |
| State 2 | -2.5984 | 0.0156^*^ |
| State 3 | 2.9302 | 0.0086^*^ |
| State 4 | 3.1496 | 0.0086^*^ |
| State 5 | -0.1705 | 0.8649 |
| State 6 | -0.2026 | 0.8649 |
| **Persistence** | **T value** | **P value (FDR adjusted)** |
| State 1 | -3.2723 | 0.0081^*^ |
| State 2 | -2.9275 | 0.0120^*^ |
| State 3 | 1.5616 | 0.1811 |
| State 4 | 1.2092 | 0.2745 |
| State 5 | -1.0259 | 0.3068 |
| State 6 | -2.3135 | 0.0444^*^ |
| **Counts** | **T value** | **P value (FDR adjusted)** |
| State 1 | -1.7997 | 0.1112 |
| State 2 | -1.4180 | 0.1902 |
| State 3 | 2.1838 | 0.0922 |
| State 4 | 3.3455 | 0.0064^*^ |
| State 5 | 0.4108 | 0.6819 |
| State 6 | 1.9082 | 0.1112 |

^*^ indicates the FDR-adjusted p-value < 0.05.

**Table S4.** The group (SZ and HC) and frequency (slow-4 and slow-5) main effects of CAP dynamics.

| **Fraction of time** | **F value** | **P value (FDR adjusted)** |
| --- | --- | --- |
| **Group** | | |
| State 1 | 9.79 | 0.0026 |
| State 2 | 13.24 | 5.30 × 10^-4^ |
| State 3 | 21.62 | 1.58 × 10^-5^ |
| State 4 | 20.87 | 2.19 × 10^-5^ |
| **Frequency** | | |
| State 4 | 7.00 | 0.0101 |
| State 5 | 6.40 | 0.0137 |
| State 6 | 18.99 | 4.54 × 10^-5^ |
| **Persistence** | **F value** | **P value (FDR adjusted)** |
| **Group** | | |
| State 2 | 5.32 | 0.0241 |
| State 3 | 10.36 | 0.0020 |
| State 4 | 15.50 | 1.97 × 10^-4^ |
| State 5 | 4.52 | 0.0371 |
| **Frequency** | | |
| State 1 | 543.69 | < 0.0001 |
| State 2 | 512.14 | < 0.0001 |
| State 3 | 764.14 | < 0.0001 |
| State 4 | 981.03 | < 0.0001 |
| State 5 | 727.54 | < 0.0001 |
| State 6 | 571.64 | < 0.0001 |
| **Counts** | **F value** | **P value (FDR adjusted)** |
| **Group** | | |
| State 1 | 7.16 | 0.0094 |
| State 2 | 5.49 | 0.0220 |
| State 3 | 12.96 | 6.00 × 10^-4^ |
| State 4 | 14.05 | 3.69 × 10^-4^ |
| **Frequency** | | |
| State 1 | 1.17 × 10^3^ | < 0.0001 |
| State 2 | 1.25 × 10^3^ | < 0.0001 |
| State 3 | 1.77 × 10^3^ | < 0.0001 |
| State 4 | 2.12 × 10^3^ | < 0.0001 |
| State 5 | 1.09 × 10^3^ | < 0.0001 |
| State 6 | 1.05 × 10^3^ | < 0.0001 |

**Table S5.** The classification results (SZ vs HC) using the temporal features.

|  | **typical range** | **slow-5** | **slow-4** | **slow-5 + slow-4** |
| --- | --- | --- | --- | --- |
| **AUC** | 0.7034 | 0.7173 | 0.6299 | **0.7356** |
| **ACC** | 0.6667 | 0.7029 | 0.5870 | **0.7391** |
| **SE** | 0.7246 | 0.7971 | 0.6957 | **0.7971** |
| **SP** | 0.6087 | 0.6087 | 0.4783 | **0.6812** |

AUC: area under curve, ACC: accuracy, SE: sensitivity, SP: specificity.

**Table S6.** The classification results (SZ vs HC) using the spatial features.

|  | **typical range** | **slow-5** | **slow-4** | **slow-5 + slow-4** |
| --- | --- | --- | --- | --- |
| **AUC** | 0.9443 | 0.9200 | 0.9477 | **0.9630** |
| **ACC** | 0.8841 | 0.8551 | 0.8841 | **0.8913** |
| **SE** | 0.8986 | 0.8696 | 0.8696 | **0.8986** |
| **SP** | 0.8696 | 0.8406 | 0.8986 | **0.8841** |

AUC: area under curve, ACC: accuracy, SE: sensitivity, SP: specificity.
